# Nonlinear microscale mechanics of actin networks governed by coupling of filament crosslinking and stabilization

**DOI:** 10.1101/2022.10.25.513787

**Authors:** Mike Dwyer, Rae M. Robertson-Anderson, Bekele J. Gurmessa

## Abstract

Actin plays a vital role in maintaining the stability and rigidity of biological cells while allowing for cell motility and shape change. The semiflexible nature of actin filaments – along with the myriad actin-binding proteins (ABPs) that serve to crosslink, bundle, and stabilize filaments – are central to this multifunctionality. The effect of ABPs on the structural and mechanical properties of actin network mechanics has been the topic of fervent investigation over the past few decades, revealing diverse structures from isotropic percolated networks to heterogeneous bundles that depend on the crosslinker type and concentration. Yet, the impact of filament stabilization and stiffening via ABPs on the nonlinear response of crosslinked networks has yet to be explored. Here, we perform optical tweezers microheology measurements to characterize the nonlinear force response and relaxation dynamics of actin networks in the presence of varying concentrations of *α*-actinin, which transiently crosslinks actin filaments, and phalloidin, which stabilizes filamentous actin and increases its persistence length. We show that crosslinking and stabilization can act both synergistically and antagonistically to tune the network resistance to nonlinear straining. For example, phalloidin-stabilization leads to enhanced elastic response and reduced dissipation at large strains and timescales, while the initial microscale force response is reduced compared to networks without phalloidin. Moreover, we find that stabilization switches this initial response from that of stress-stiffening to softening despite the increased filament stiffness that phalloidin confers. Finally, we show that both crosslinking and stabilization are necessary to elicit these emergent features, while the effect of stabilization on networks without crosslinkers is much more subdued. We suggest that these intriguing mechanical properties arise from the competition and cooperation between filament connectivity, bundling, and rigidification, shedding light on how ABPs with distinct roles can act in concert to mediate diverse mechanical properties of the cytoskeleton and bio-inspired polymeric materials.

## 1. Introduction

The mechanical properties of networks of actin filaments have been the topic of fervent investigation for decades due to the key roles they play in maintaining the stability and rigidity of biological cells while also allowing for cell motility and shape change [1–5]. This multifunctionality is enabled by the semiflexible nature of actin filaments and the myriad actin-binding proteins (ABP) that can, e.g., crosslink, stabilize, stiffen, and bundle actin filaments, resulting in viscoelastic networks with wide-ranging mechanical properties [2,6–11]. A notable feature of these networks, relevant to both cellular biology and polymer physics, is their unique nonlinear response to strain and the roles that filament stiffness, crosslinking, and concentration play in this signature nonlinear response [12–15].

In the absence of ABPs, actin filaments above a critical concentration form entangled networks in vitro with linear viscoelastic properties that can be described reasonably well by reptation-based models for entangled semiflexible polymers [16,17]. The nonlinear response of entangled actin networks is more complex and has been shown to exhibit varying degrees of stress stiffening and softening depending on the spatiotemporal scale of the strain and the network concentration [12,13,18]. Forced disentanglement, shear-thinning, entropic stretching, and strain alignment have also all been implicated as important features of the nonlinear response of entangled actin networks [12,13,19,20].

Introducing crosslinking ABPs into entangled actin networks greatly enhances the elastic contribution to the mechanical response by suppressing thermal fluctuations and disentanglement. However, dissipative processes still contribute appreciably to the mechanics at low ABP:actin ratios *R* and for transient crosslinkers such as *α*-actinin, which continuously bind and unbind to actin filaments to allow for network rearrangement [21]. Even in the case of static crosslinking, such as in studies that use avidin proteins to bind biotinylated actin filaments, dissipation, relaxation, and plastic rearrangement have been reported to contribute to the response to nonlinear straining due to forced rupturing of crosslinker bonds [12,22–24]. The type and concentration of crosslinking ABPs also control the extent to which ABPs isotropically crosslink actin filaments to form a homogeneous well-connected network of individual filaments or bundle filaments to form a more loosely connected and heterogeneous network of stiff fibers (i.e., multi-filament bundles). These alterations to network connectivity and fiber stiffness play important roles in the diverse mechanical and structural features that have been reported for actin networks crosslinked by a range of ABPs including, e.g., *α*-actinin, scruin, fascin, and filamin [13,14,25–36].

While bundling via ABPs increases the stiffness of actin network fibers [29,34,37–42], stabilizing ABPs such as phalloidin stiffen individual filaments [43–46]. Phalloidin selectively binds to filamentous actin to suppress treadmilling [44,47], thereby accelerating the polymerization rate and lowering the critical polymerization concentration [44,47]. Phalloidin-stabilized filaments have been reported to have a persistence length of *l_p_* ≃ 17 *μ*m, which is nearly twice *l_p_* ≃ 10 *μ*m measured for unstabilized actin filaments [46,48–50]. Moreover, stabilized filaments have been reported to form bundles similar to those formed by high concentrations of crosslinking ABPs [33]. Filament bundling contributes to the viscoelastic response of actin networks in a manner highly dependent on network concentration due to competing effects of increased filament rigidity and reduced network connectivity [7,18,33,51–54].

Here, we investigate the coupled roles of stabilization, crosslinking, and entanglement density on the microscale nonlinear response of in vitro actin networks. Specifically, we use optical tweezers microrheology to measure the nonlinear force response and relaxation dynamics of entangled actin networks with phalloidin:actin ratios of *R_p_* = 0 − 0.02, *α*-actinin:actin ratios of *R_α_* = −0 0.03, and actin concentrations of *c_a_* = 2.9 − 11.6 *μ*M (Fig 1). We find that while phalloidin-stabilization increases the elastic response and stiffness at mesoscopic scales; it surprisingly suppresses stress-stiffening and lowers the resistive force that crosslinked networks exhibit at the microscale. We also uncover intriguing nonmonotonic dependence of stiffness and relaxation dynamics on *R_p_*, *R_α_* and *c_a_*, which we rationalize as arising from the distinct roles that filament bundling and connectivity play in the nonlinear response of actin networks.

**Figure 1.**
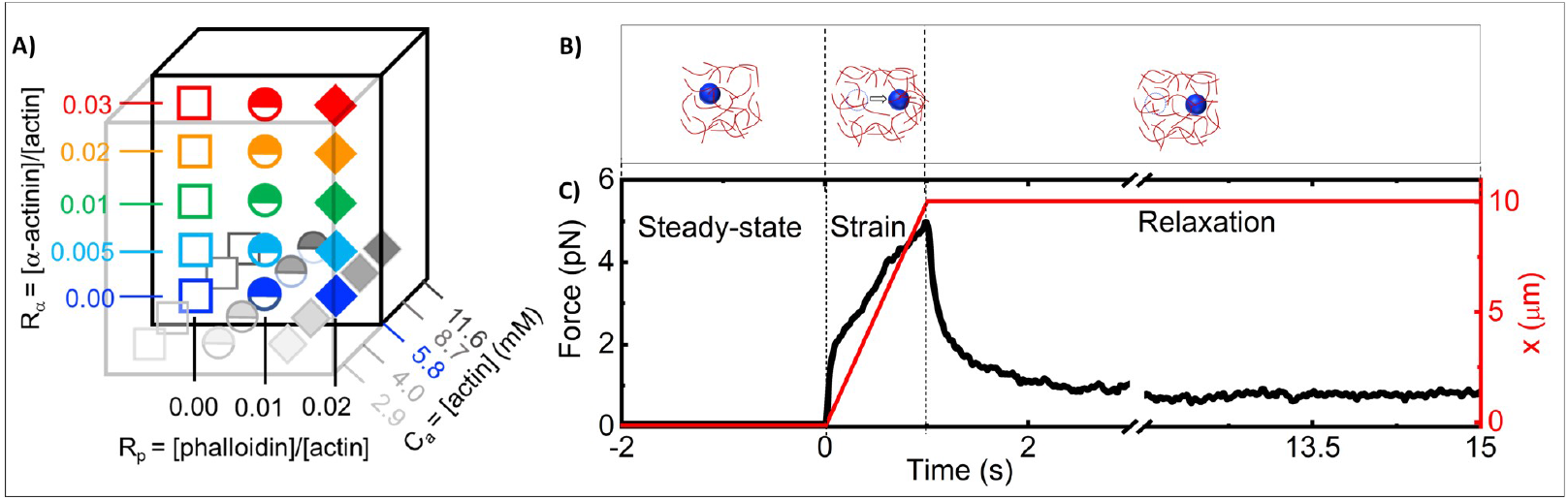
Optical tweezers microrheology characterizes the nonlinear force response of actin networks with varying concentrations of actin, *α*-actinin and phalloidin: A) Cartoon of experimental parameter space of actin networks with varying: *α*-actinin:actin molar ratios, *R_α_* = 0–0.03 (*y*-axis), phalloidin:actin molar ratios *R_p_* = 0–0.02 (*x*-axis), and molar actin concentrations, *c_a_* = 2.9–11.6 *μ*M (*z*-axis). (B) Optical tweezers microrheology (OTM) measurement protocol: an optically trapped 4.2-*μ*m diameter microsphere probe (blue) embedded in an actin network (red) is: held fixed for 15 s to allow the network and probe to reach steady-state and establish a baseline (left, steady-state), displaced *x* = 10 *μ*m at a constant speed *v* = 10 *μ*m/s by moving the piezoelectric stage relative to the trap (middle, strain), and held fixed while the network relaxes to a new steady-state (right, relaxation). The dashed circles represent the initial probe position, and the arrow indicates the trap motion during strain. (C) A sample OTM measurement showing the measured force exerted on the trapped probe (black) and the stage position versus acquisition time. The dashed vertical lines divide the data into the three phases depicted in (B).

## 2. Materials and Methods

### Proteins

Rabbit skeletal actin, *α*-actinin, and Acti-Stain 555 Phalloidin (Cytoskeleton, Inc) were reconstituted to 46 *μ*M, 10 *μ*M and 14 *μ*M and stored, respectively, at −80 °*C* in G-buffer [2 mM Tris pH 8.0, 0.5 mM DTT, 0.1 mM CaCl_2_, 0.2 mM ATP], at −80 °*C* in [4 mM Tris-HCl pH 7.6, 4 mM NaCl, 20 μM EDTA, 1% (w/v) sucrose, 0.2% (w/v) dextran], and at −20 °*C* in 100% methanol. Alexa-568 labeled actin (Thermo Fisher Scientific) was reconstituted to 35 *μ*M and stored at −80 °*C* in G-buffer [2 mM Tris pH 8.0, 0.5 mM DTT, 0.1 mM CaCl_2_].

### Network Formation

We mix unlabeled G-actin and either (i) Alexa568-actin or (ii) Acti-Stain-555-phalloidin with oxygen scavenging agents (4.5 *μ*g/ml glucose, 0.005% *β*-mercaptoethanol, 4.3 *μ*g/ml glucose oxidase, 0.7 *μ*g/ml catalase) to final actin concentrations of 2.9–11.6 *μ*M in F-buffer [10 mM Imidazole pH 7.0, 50 mM KCl, 1 mM MgCl_2_, 1 mM EGTA, 0.2 mM ATP]. We include 568-actin or 555-phalloidin at (i) a 1:10 ratio of labeled:unlabeled actin or (ii) phalloidin:actin ratios of *R_p_* = 0.01 and 0.02. We also include *α*-actinin at molar *α*-actinin:actin ratios of *R_α_* = 0, 0.005, 0.01, 0.02, and 0.03 (Fig 1A), which have been reported to result in isotropic, crosslinked networks with small-scale bundles forming above *R_α_* ≃ 0.02 [21,33].

We immediately flow the solution into a sample chamber comprised of double-sided tape as a spacer between a glass slide and coverslip, seal the chamber with epoxy, and allow the actin to polymerize at room temperature for 1 hr. For all networks, we add a trace of 4.2-*μ*m diameter polystyrene microspheres (Bangs Laboratory, Inc.) prior to polymerization to allow for microrheology measurements (Fig 1B). The predicted mesh sizes of the actin networks at the concentrations we examine span 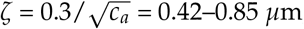, where *c_a_* is the actin concentration in mg/mL [55].

### Microscopy

To characterize the structure of the different actin networks, we use a Leica TCS SP8 laser scanning confocal microscope with a 63x 1.4 NA objective and 561 nm laser to image fluorescently labeled filaments in actin networks. We collect z-stacks of 21 images of 512 × 512 pixels, each with a z-depth of 0.5 *μ*m. The resulting z-stacks are 123 × 123 × 10.5 *μ*m.

### Microrheology

We used an optical trap built around an IX73 fluorescence microscope (Olympus, Melville, NY) with a 1064 nm Nd:YAG fiber laser (BKtel, RPMC Lasers, Inc) focused with a 60x 1.42 NA oil immersion objective (UPLXAPO60XO, Olympus). A position-sensing detector (PSM2-10Q PSD/OT-301, ON-TRAK Photonics, Inc) measured the deflection of the trapping laser, which is proportional to the force acting on the trapped probe over our entire force range. The trap stiffness was calibrated via Stokes drag in water [56] and passive equipartition methods [57]. During measurements, a probe embedded in the network is trapped and moved 10 *μ*m at a constant speed of 10 *μ*m/s relative to the sample chamber via steering of a nanopositioning piezoelectric stage (PDQ-250, Mad City Laboratories) while measuring both the laser deflection and stage position at a rate of 20 *kHz* during the three phases of the experiment: equilibration (5 *s*), strain (1 *s*) and relaxation (15 *s*) (Fig 1B). The strain rate was chosen to be higher than the previously determined strain rate necessary for the onset of nonlinear mechanics [58]. Displayed force curves are averages of 30 trials using 30 different probes at different locations in the sample chamber. Custom-written LabVIEW code was used to perform measurements and acquire data, while data analysis was carried out with custom-written Matlab programs.

## 3. Results and Discussion

We first aim to characterize the effect of phalloidin-stabilization on the nonlinear force response and relaxation dynamics and steady-state structure of actin networks crosslinked with *α*-actinin.

### Network structure

To verify that the actin networks we examine follow the previously reported structural trends [33], and to shed light on the structural impact of phalloidin-stabilization, we acquire three-dimensional fluorescence confocal image stacks of the labeled actin networks. Fig 2A,B, which shows *z*-projections of images, color-coded by *z*-height, and single images and zoom-ins to the right of each colorized projection, displays network structure without (Fig 2A, *R_p_* = 0) and with (Fig 2B, *R_p_* = 0.02) phalloidin for increasing crosslinker concentrations *R_α_* (left to right). Without phalloidin, there is a stark difference between networks with and without crosslinkers, with the unlinked network (*R_α_* = 0) appearing much more homogeneous and densely filled with individual filaments (Fig 2A). The lack of clear discernible filaments indicates enhanced Brownian noise compared to the crosslinked cases, as filaments can fluctuate over the course of the acquisition. Conversely, the crisp, high-contrast fibers seen in crosslinked networks (Fig 2B), coupled with larger visible dark regions void of filaments, indicate that the crosslinked fibers are highly rigid and static over the course of acquisition and that the fibers are likely small bundles rather than single filaments.

**Figure 2.**
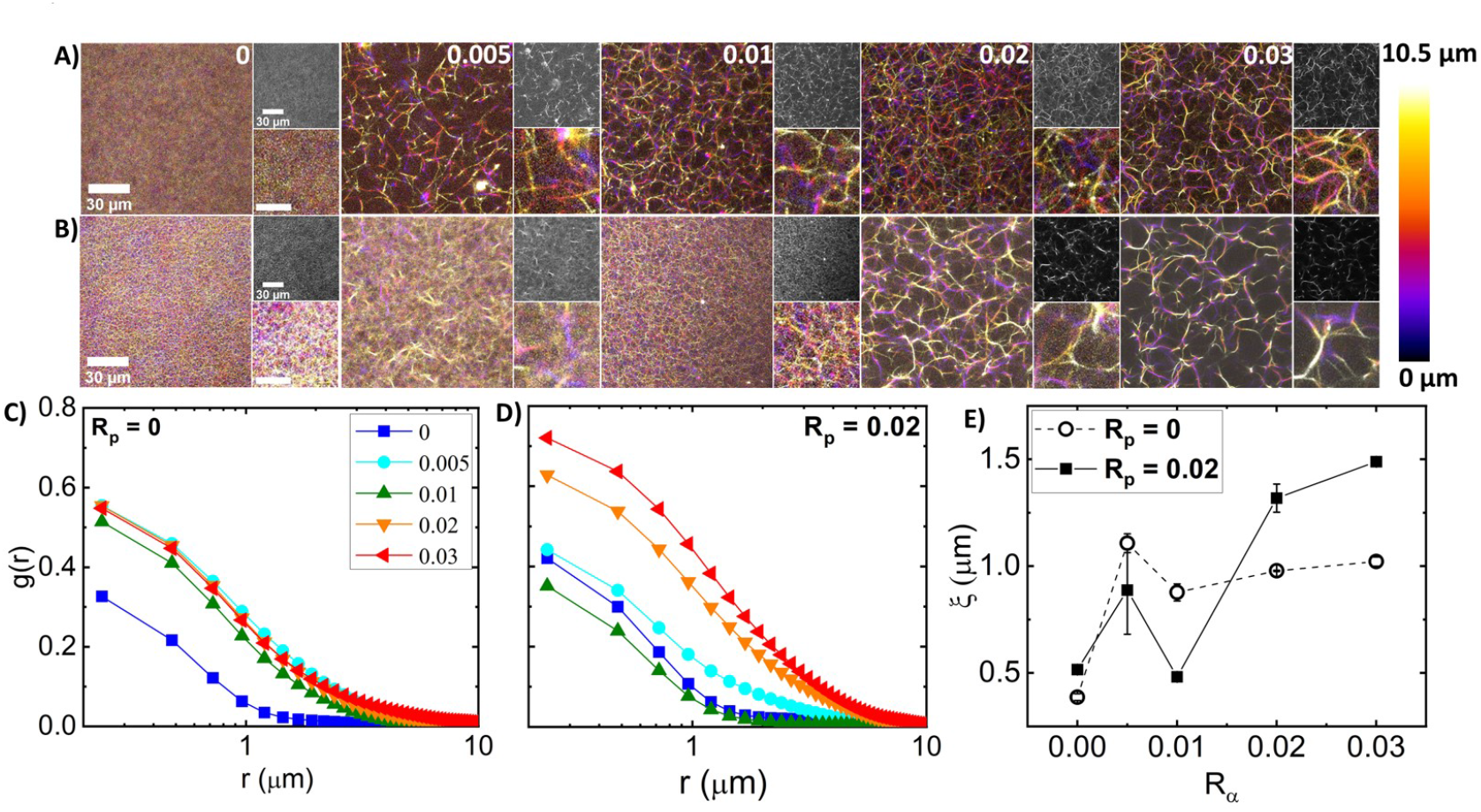
The structure and mobility of actin networks depend on the degree of filament crosslinking and stabilization. (A, B) Laser scanning confocal microscopy imaging of 5.8 *μ*M fluorescent-labeled actin networks with varying *α*-actinin:actin molar ratios *R_α_* = 0–0.03, increasing from left to right as indicated at the top right corner of each image. Networks (A) without phalloidin (*R_p_* = 0) and (B) with phalloidin:actin molar ratio *R_p_* = 0.02 depict the effect of filament stabilization on network structure. For each [*R_α_*,*R_p_*] combination, 3-dimensional stacks of 21 images, each of 0.5 *μ*m thickness, are captured using a Leica TCS SP8 laser scanning confocal microscope with 60x 1.4 NA objective. Large left-hand images are color-coded *z*-projections of the image stacks, with the color scale shown to the right and scale bars representing 30 *μ*m. Small images are single slices from the stack (top) and zoom-ins of the *z*-projections (bottom) with a scale bar indicating 10 *μ*m. (C,D) Spatial image autocorrelation functions, *g*(*r*) versus radial distance *r*, computed from each image of each *z*-stack shown in (A, B), quantify the average feature size of images. The individual curves are the average *g*(*r*) values for each *R_α_*, as denoted in the legend, for networks with (C) *R_p_* = 0 and (D) *R_p_* = 0.02. (E) The correlation lengths *ξ* as a function of *R_α_*, obtained by fitting each *g*(*r*) curve shown in (C) (open symbols, dashed connecting lines) and (D) (closed symbols, solid connecting lines) to an exponentially decaying function: *g*(*r*) = *g*(0) exp (−*r*/*ξ*). Error bars are the standard error across *ξ* values measured in each image of the corresponding *z*-stack

To quantify these structural variations, we compute the spatial image autocorrelation function *g*(*r*), where *r* is the radial distance between two pixels for each image (shown in greyscale in Fig 2) in the image stack. The autocorrelation *g*(*r*) is a measure of how correlated the intensities of two pixels a given distance *r* away from one another are. The distance *r* over which *g*(*r*) decays, often termed the correlation length *ξ*, indicates the average size of features in the image. For an isotropic fiber network, *ξ* is comparable to the mesh size. As shown in Fig 2C, the autocorrelation is lower for all *r* values for the unlinked network, indicating minimal spatial structure and increased thermal noise compared to the crosslinked networks. Introducing crosslinkers significantly increases the structural correlation for all radial distances and also increases the lengthscale over which *g*(*r*) decays, both trends indicative of larger and more pronounced structural features. By fitting each autocorrelation function to an exponential decay function, *g*(*r*) = *g*(0) exp (−*r*/*ξ*), we extract the correlation length, *ξ*, which corroborates our qualitative description. Namely *ξ* increases ~2-fold upon crosslinking, but remains roughly constant for all *R_α_* > 0 values (Fig 2E).

Interestingly, adding phalloidin to the networks shown in Fig 2(A) appears to either inhibit *or* promote the restructuring that *α*-actinin crosslinking induces, depending on *R_α_*. Specifically, for *R_α_* ≤ 0.01, phalloidin stabilization appears to have little impact on the apparent density or homogeneity of the networks, as seen by the similar *g*(*r*) curves and *ξ* values for *R_α_* = 0, 0.005 and 0.01 (Fig 2D,E). In contrast, in the absence of phalloidin there is evident small-scale bundling for *R_α_* = 0.005 and 0.01 networks, as shown by the larger *g*(*r*) and *ξ* values compared to the *R_α_* = 0 network (Fig 2C,E). On the other hand, for *R_α_* > 0.01, phalloidin stabilization seems to increase *α*-actinin mediated bundling, as indicated by the thicker and brighter fibers and larger empty voids in the last two columns of Fig 2(B) compared to Fig 2(A). This increased bundling is evidenced as larger *g*(*r*) values in Fig 2D compared to Fig 2C for *R_α_* > 0.01 (i.e. red and orange curves), and larger *ξ* values of phalloidin-stabilized (*R_p_* = 0.02) versus non-stabilized (*R_p_* = 0) networks for *R_α_* > 0.01 in Fig 2E.

### Nonlinear Stress Response

To determine how the seemingly antagonistic and synergistic effects of stabilization and crosslinking, suggested in Fig 2, impact the nonlinear force response of the networks, we perform optical tweezers microrheology measurements described in Methods and Fig 1. In brief, we pull an optically trapped probe through a distance *x* = 10 *μ*m at a constant speed of *v* = 10 *μ*m/s through the network and measure the force the network exerts to resist the strain. Following the strain, we hold the probe fixed at the final strain position and measure how the force decays over time. For reference, in a purely elastic material, the force *F*(*x*) would increase linearly with strain distance *x*, with the slope *dF*/*dx* indicating the effective spring constant or stiffness, while a viscous fluid would nearly instantly reach an *x*-independent plateau. Following the strain, an elastic material would retain the induced force indefinitely, while for a viscous fluid, the force would drop to zero immediately upon halting the probe motion. Entangled and crosslinked actin networks have been shown to exhibit viscoelastic features intermediate to these extremes, depending on the actin and crosslinker concentration [18,26,51,52].

Figure 3(A) shows average force response curves for networks of varying *R_α_* values, with each panel showing networks with a fixed phalloidin:actin ratio: *R_p_* = 0 (left), *R_p_* = 0.01 (middle), and *R_p_* = 0.02 (right). As shown, the force *F*(*x*) generally increases with increasing *R_α_* for all cases; however, the functional dependence on *R_α_* is distinct for different phalloidin concentrations. Moreover, phalloidin stabilization substantially increases the resistive force compared to *R_p_* = 0 for all crosslinker ratios. We quantify these trends by examining the terminal force *F_t_* reached at the end of the strain as a function of *R_α_* (Fig 3B). We observe that, without phalloidin, *F_t_* increases ~2-fold at *R_α_* = 0.01 but remains nearly constant as *R_α_* is increased further, similar to the insensitivity of the structure to *R_α_* (Fig 2). Conversely, for *R_α_* = 0.02, the terminal force, which is substantially larger than *F_t_* for *R_α_* = 0, increases monotonically with *R_α_*. Intriguingly, the reduced stabilization case (*R_p_* = 0.01) exhibits a non-monotonic dependence of *F_t_* on *R_α_*, transitioning from a value close that for *R_p_* = 0 in the absence of crosslinking to values that are higher than the *R_p_* = 0.02 network for *R_α_* = 0.005–0.02, after which *F_t_* drops modestly.

**Figure 3.**
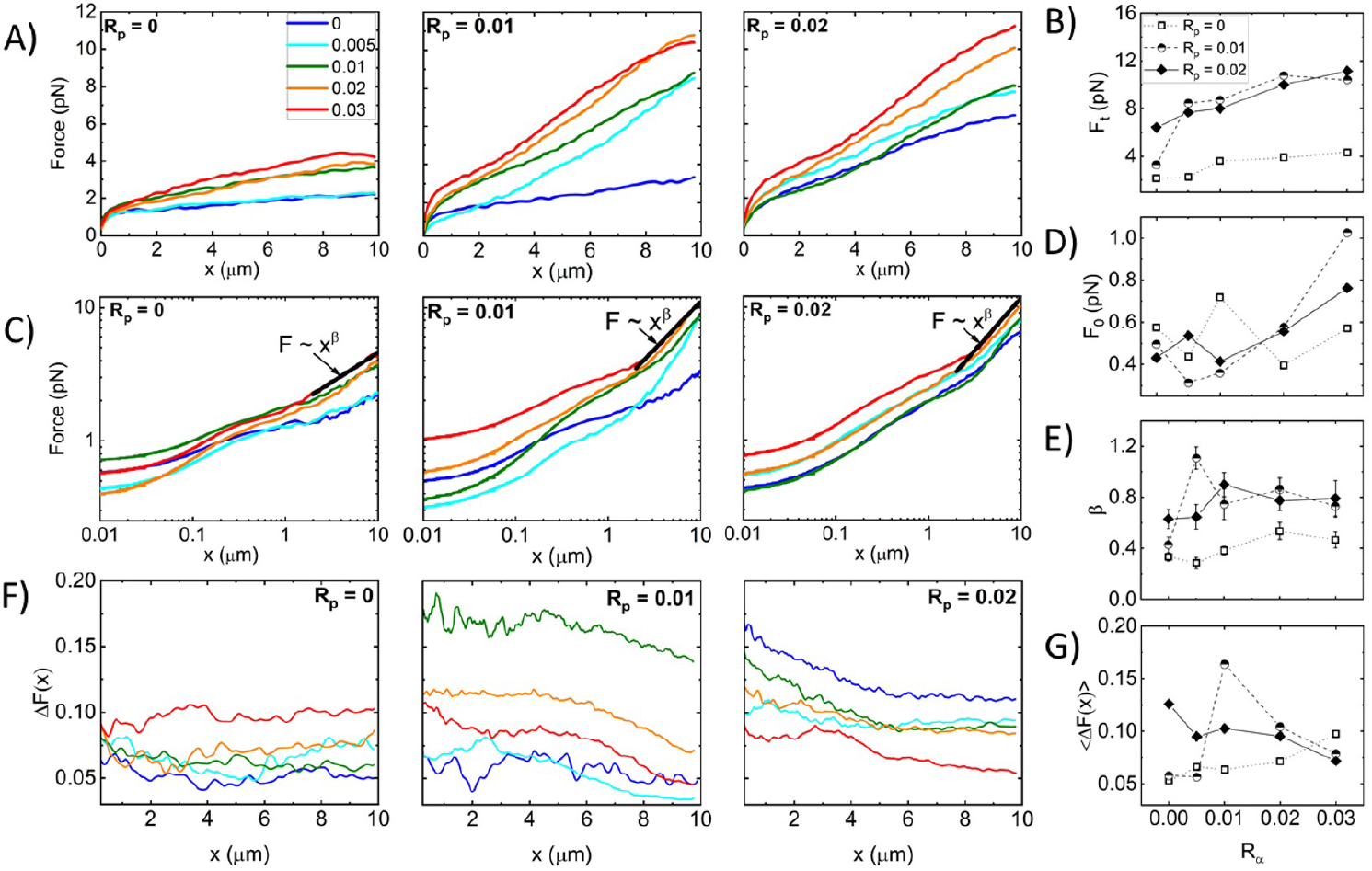
Synergistic effects of crosslinking and stabilization enable sustained elastic response of actin networks to nonlinear straining. (A) Force *F*(*x*) versus stage position *x* measured for actin networks subject to nonlinear straining (see Fig 1). Different curves in each panel correspond to *α*-actinin:actin molar ratios of *R_α_* = 0 − 0.03, color-coded according to the legend in A. Different panels display data for phalloidin:actin molar ratios of *R_p_* = 0 (left), *R_p_* = 0.01 (middle), and *R_p_* = 0.02 (right). (B) Terminal force reached at the end of the strain *F_t_* as a function of *R_α_* for *R_p_* = 0 (open squares, dotted connecting lines), *R_p_* = 0.01 (half-filled circles, dashed connecting lines), and *R_p_* = 0.02 (solid diamonds, solid connecting lines). (C) Data shown in (A) plotted on a log-log scale to highlight the trends seen for the initial force *F*(*x* = 0) = *F*_0_ and power-law scaling of *F*(*x*) near the end of the strain (*x* ≈ 1). Fitting the large strain data to a power-law *F*(*x*) ~ *x^β^* yields the scaling exponent *β* that quantifies the degree of elastic storage maintained at the end of the strain. (D) Initial force measured at the beginning of the strain *F*_0_ and (E) power-law scaling exponent *β* determined from fits to *F*(*x*) ~ *x^β^* depicted in B versus *R_α_* for *R_p_* = 0 (open squares, dotted connecting lines), *R_p_* = 0.01 (half-filled circles, dashed connecting lines), and *R_p_* = 0.02 (solid diamonds, solid connecting lines. (F) Fractional spread in force Δ*F*(*x*) for each position *x* and each condition, determined by computing the standard error across 30 individual trials and normalizing by the average value plotted in A: Δ*F*(*x*) = *SE*_*F*(*x*_)/⟨*F*(*x*)⟩. (G) Δ*F*(*x*) averaged over the strain position *x*, resulting in a single value for each curve shown in (C) with error bars denoting the standard error across *x* values.

To shed light on the mechanisms underlying these complex trends, we note that the functional forms of the *F*(*x*) curves differ for different crosslinker and phalloidin concentrations, such that the initial force *F*_0_ may exhibit different dependence on *R_α_* and *R_p_* than the terminal force *F_t_*. To better visualize the initial force response and the dependence of *F* on *x*, we plot the data shown in Fig 3A on a log-log scale (Fig 3C), from which we observe that indeed the small strain response displays a more complex dependence on *R_α_* and *R_p_* as quantified in Fig 3D. Without crosslinkers (i.e., *R_α_* = 0), phalloidin stabilization actually reduces *F*_0_, whereas for *R_p_* > 0.01, stabilization increases *F*_0_ (Fig 3D). A similar phenomenon has been reported for entangled composites of actin filaments, and rigid microtubules, whereby composites with more microtubules compared to actin exhibited lower *F*_0_ values compared to actin-rich composites, but this trend flipped at larger lengthscales where the resistive force was substantially higher for microtubule-rich composites [59]. This work showed that the increased initial force for actin-rich composites arose from the decreased mesh size and increased flexibility compared to microtubule-rich composites, which, in turn, increased the poroelastic contributions to the initial stress response [59]. Conversely, the lower terminal force *F_t_* arose from the semiflexible actin filaments being able to more readily dissipate induced stress compared to their rigid microtubule counterparts. Similarly, we may understand the reduced initial force and increased terminal force for phalloidin stabilized filaments as arising from stiffening and bundling of filaments, making them more akin to microtubules, compared to actin networks without phalloidin.

Fig 3C also reveals a discrete shift at *x* ≃ 1 *μ*m to a steeper power-law dependence of the force on strain distance, *F*(*x*) ~ *x^β^*, which is more pronounced for phalloidin-stabilized networks. In other words, rather than softening to a viscous-dominant regime in which *β* ~ 0, as has been previously reported for both entangled and crosslinked actin networks [44–46], phalloidin-stabilized networks appear to stiffen at large strains, reaching scaling close to the elastic limit *β* ~ 1(Fig 3E). Moreover, the scaling exponent displays a nonmonotonic dependence on crosslinking density, with the most pronounced stiffening occurring for 0 < *R_α_* < 0.03.

Our results suggest that both fiber stiffness and mesh size play important roles in the force response, which we also expect to affect the heterogeneity between trials measured in different regions of the sample and across different samples. Because the probe diameter of *a* = 4.2 *μ*m is an order of magnitude larger than the predicted mesh size (*ζ* ≃ 0.6 *μ*m) and the measured correlation length *ξ* of the entangled actin network, the probe senses an effectively homogeneous continuum as it moves through the [*R_α_* = 0,*R_p_* = 0] network. However, if the mesh size increases significantly due to bundling, as indicated in Fig 2 for networks with high *R_α_* and *R_p_* > 0, then the probe may detect these structural heterogeneities as it moves through the network. Moreover, while the semiflexibility of individual actin filaments allows them to bend, stretch and reorient during the strain such that the bead does not encounter discrete rigid entities, this assumption does not necessarily hold for networks of rigid fibers [59] or highly crosslinked polymers [60,61].

To quantify this heterogeneity we compute the fractional spread in force values Δ*F*(*x*) = *SE_f_* /*F*(*x*) that contribute to each *F*(*x*) curve shown in Fig 3A,B, where *SE_f_* is the standard error across the 30 individual force values *f* (*x*) measured at each *x* for each condition (Fig 3F). As shown, without phalloidin stabilization, Δ*F*(*x*) is largely independent of strain distance and unaffected by *R_α_* until *R_α_* > 0.01 in which Δ*F*(*x*) increases modestly. Phalloidin stabilization leads to generally larger fractional spreads than without stabilization (*R_p_* = 0) for all crosslinker ratios and decreases with increasing strain distance. These general trends can be seen in Fig 3G, which also shows that the *x*-averaged fractional force spread 〈Δ*F*(*x*)〉 displays an emergent non-monotonic dependence on *R_α_*, with a maximum at *R_α_* = 0.005 for the more weakly stabilized networks (*R_p_* = 0.01). The *x*-dependence suggests that the network is heterogeneous on microscales that tend toward homogeneity at scales larger than ~4 *μ*m. Substantial bundling and increased stiffness of fibers, without strong connectivity between fibers, may contribute to this effect, which we explore further below.

### Time-varying Network Stiffness during Strain

Figure 3 shows that all networks exhibit varying degrees of elastic stiffness and viscous dissipation that depend not only on the network composition (i.e., *R_α_* and *R_p_*) but also on the strain distance *x*. To quantify these nonlinear stress characteristics, we evaluate the effective differential modulus *K* = *dF*/*dx*, which quantifies the elasticity or stiffness of the network as a function of time *t* = *x*/*v* during the strain, with a purely viscous response yielding *K* ≈ 0 (Fig 4).

**Figure 4.**
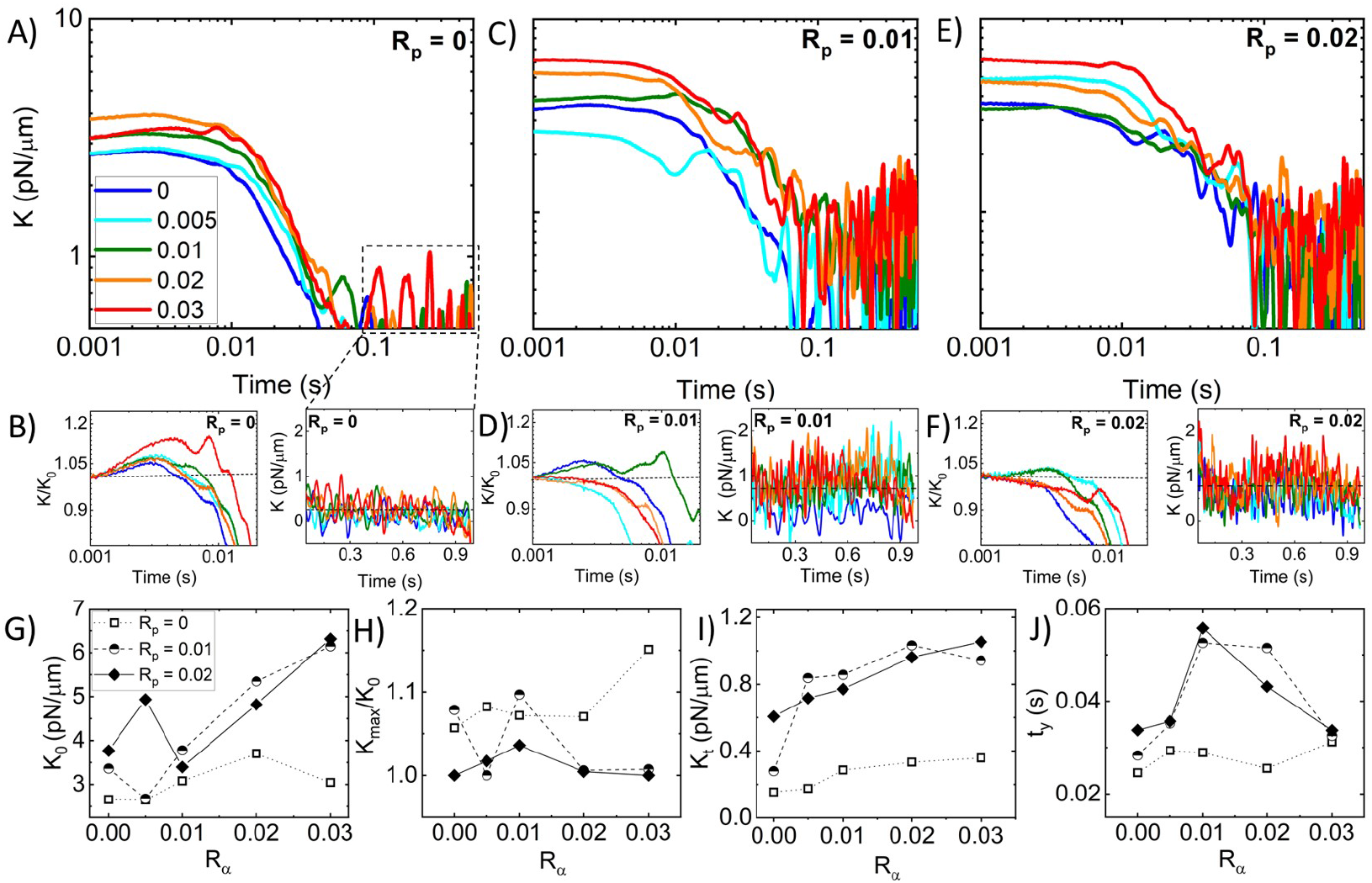
Stabilization suppresses the stress-stiffening behavior of crosslinked networks while increasing the response stiffness and associated softening timescales. (A) Effective differential modulus *K*(*t*) = *dK*(*x*, *t*)/*dx* as a function of time during strain *t* = *v*/*x*, computed from the *R_p_* = 0 data shown in 3A. The color-coded curves are for *R_α_* values indicated in the legend. (B) *K*(*t*) data shown in (A) normalized by the corresponding initial value *K*_0_ (left) with the horizontal dashed line at *K*(*t*)/*K*_0_ = 1 guiding the eye to show stress stiffening (*K*(*t*)/*K*_0_ > 1) or softening (*K*(*t*)/*K*_0_ < 1). Zoom-ins of *K*(*t*) near the end of the strain (right) where *K*(*t*) is approximately constant, with the dashed horizontal line denoting the average. (C-F) Metrics plotted in A and B for networks with *R_p_* = 0.01 (C,D) and *R_p_* = 0.02 (E,F). (G-J) Quantities computed from the data shown in A-F as functions of *R_α_* for *R_p_* = 0 (open squares, dotted connecting lines), *R_p_* = 0.01 (half-filled circles, dashed connecting lines), and *R_p_* = 0.02 (solid diamonds, solid connecting lines): (G) Initial differential modulus *K*_0_; (H) Degree of stress-stiffening, quantified as *K_max_* /*K*_0_ and equal to 1 for networks which only display softening (i.e.*K_max_* = *K*_0_); (I) Terminal stiffness *Kt* computed by averaging over the *K*(*t*) data shown in the right-hand panels of B,D,F; (J) Softening time *ty*, defined as the time at which *K*(*t*) = *K*_0_/*e*

Previous studies have shown that statically crosslinked actin networks subject to nonlinear straining exhibited initial stress stiffening, i.e., increasing *K*, followed by softening (decreasing *K*) to an *x*-independent regime [22,26,62]. Without phalloidin stabilization (Fig 4(A)), we see similar behavior, with all networks exhibiting initial stiffness *K*_0_ and stress stiffening *K*(*t*)/*K*_0_ > 0 that is generally more pronounced at higher *R_α_* values ((Fig 4B, left). After stiffening to a maximum value *K_max_*, networks soften to a nearly viscous terminal regime with small stiffness values *K_t_* that are largely independent of *x* ((Fig 4B, right).

Adding phalloidin to the networks increases the initial and terminal stiffness values, *K*_0_ and *K_t_* (Figs 4C-F), as we might expect given the increased elastic contributions to the force response seen in Fig 3 and the increased filament rigidity that phalloidin confers. However, quite unexpectedly, phalloidin stabilization suppresses the stress-stiffening behaviors seen in Fig 4(B), such that the majority of networks for both phalloidin concentrations exhibit purely softening behavior (Fig 4D,F). Moreover, this suppression is stronger for higher *R_α_* values and for the higher phalloidin concentration *R_p_* = 0.02. Stress stiffening is typically associated with affine stretching and alignment with the strain, as opposed to non-affine bending and dissipative fluctuations, which are most prevalent in the response of crosslinked networks and entangled networks subject to nonlinear straining [18,26,59]. As such, the softening of phalloidin-stabilized networks, most pronounced at higher *R_α_* values, suggests enhanced bundling as compared to networks with lower *R_α_* values and no stabilization, as seen in Fig 2, which comes at the cost of strong network connectivity necessary for affine stretching and deformation.

These complex effects can be seen more clearly in Fig 4(G–J) in which the initial stiffness *K*_0_, the relative stiffening *K_max_*/*K*_0_, and the terminal stiffness *K_t_* for all three *R_p_* values are plotted versus *R_α_*. We find that *K*_0_ is not only larger for phalloidin-stabilized networks compared to the *R_p_* = 0 case, but the increase with increasing crosslinking is substantially steeper. Conversely, we see nearly the opposite trend for the relative stiffening in which the *R_p_* = 0 network generally has the highest degree of stiffening, whereas most of the values for the stabilized networks exhibit the floor value of *K_max_*/*K*_0_ = 1, indicative of softening. Despite the lack of stiffening, phalloidin-stabilized networks retain substantially more elastic stiffness in the steady-state (*t*-independent) region of the stress response compared to the non-stabilized network. Taken together, our results suggest that the increased rigidity of the phalloidin-stabilized bundles causes an immediate elastic response as their rigid-rod-like conformation inhibits entropic stretching (required for stiffening) and dissipative bending and fluctuations (required for *K_t_* → 0).

We expect the competition between rigidification and connectivity to non-trivially impact the timescales required for the network to yield or relax to a steady-state stiffness, which we quantify as the time *t_y_* at which *K* reaches *K* = *K*_0_/*e*. Fig 4(J) reveals that without stabilization the yielding time is largely unaffected by the degree of crosslinking, whereas both stabilized networks exhibit a stark nonmonotonic dependence on *R_α_*, with the maximum *t_y_* value reached at *R_α_* = 0.01. Moreover, *t_y_* values for stabilized networks are nearly identical to the non-stabilized network at the highest crosslinker density (*R_α_* = 0.03) and in the absence of crosslinking (*R_α_* = 0). This trend suggests that the stiffness of the individual filaments may not affect the yield time as much as the propensity to crosslink and bundle. In the following section, we explore this interpretation and more closely investigate the impact of *R_α_* and *R_p_* on network relaxation dynamics.

### Nonlinear Relaxation Dynamics

As described in Fig 1(B, C), following the 10 *μ*m strain, we hold the bead fixed at this maximum strain position and measure the relaxation of the induced force over time. Recall that purely elastic systems maintain the induced force indefinitely (no relaxation), while viscous systems exhibit immediate and complete dissipation. Fig 5(A-C), which displays the relaxation profiles that follow the force response curves shown in Fig 3, shows that all networks undergo some degree of relaxation while also maintaining some residual force *F_R_* at the end of the relaxation period. In most cases, *F_R_* increases with increasing *R_α_* and *R_p_*, as shown in Fig 5(D), indicating that crosslinking and stabilization both contribute to enhancing elastic storage, with crosslinking having a more significant impact. As in previous studies on crosslinked and bundled actin networks [26, 59], we find that, for all networks, the relaxation of the force to the residual plateau *F_R_* can be described well by a sum of two decaying exponentials with a long-time residual: *F*(*t*) = *C*_1_ exp(−*t*/*τ*_1_) + *C*_2_ exp (−*t*/*τ*_2_) + *F_R_*. The two distinct characteristic decay times *τ*_1_ and *τ*_2_, determined from the fits of the data to this function, are measures of the timescales associated with two independent relaxation mechanisms. The corresponding coefficients *C*_1_ and *C*_2_ are measures of the relative contributions of each mechanism to the overall relaxation.

**Figure 5.**
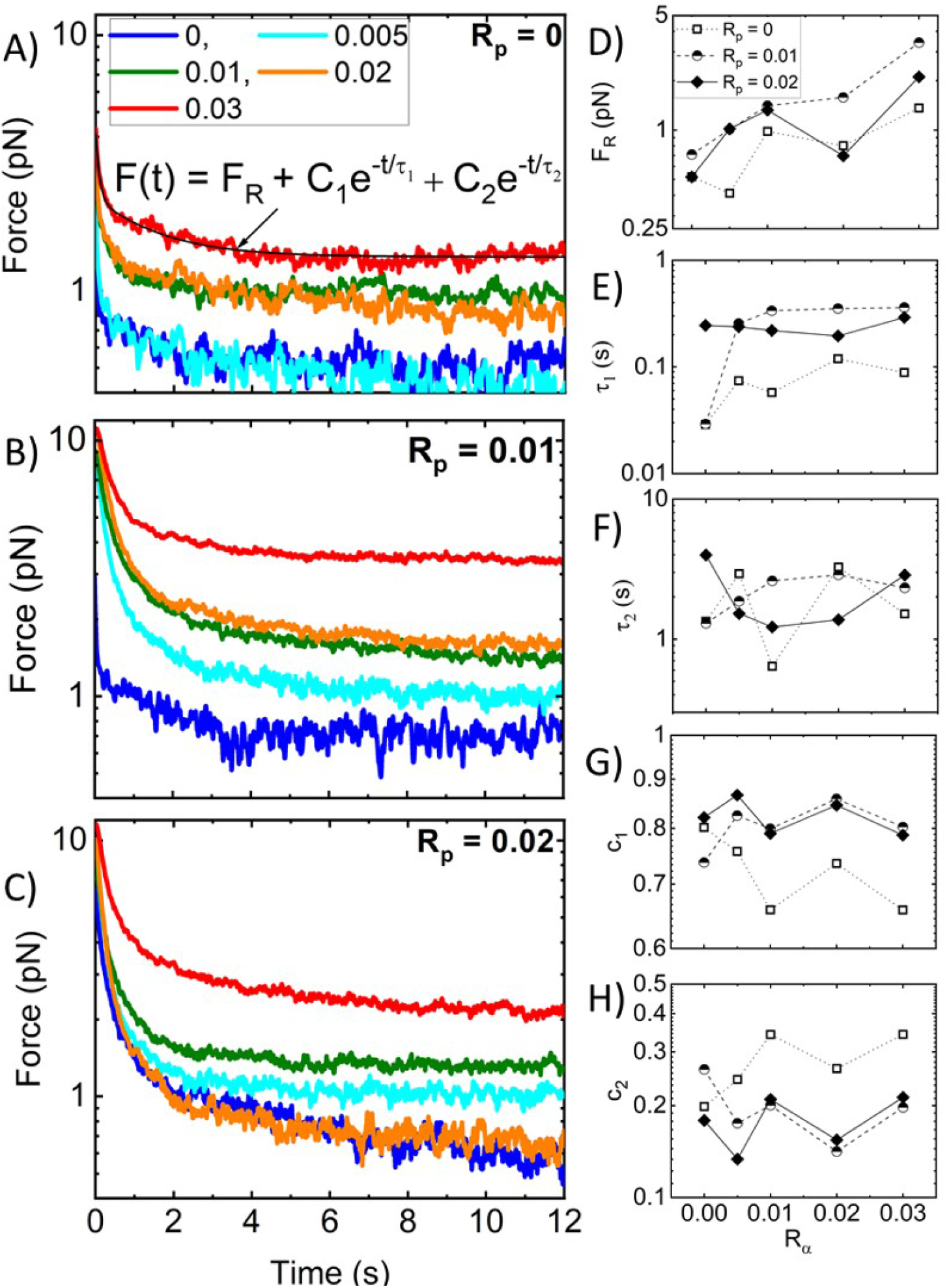
Strain-induced force exponentially relaxes over time to a residual value *F_r_* dependent on *R_p_* and *R_α_*. (A-C) Time-dependent relaxation of force *F*(*t*) following strain for actin networks with varying *R_α_* values, color-coded according to the legend, and with *R_p_* values of (A) 0, (B) 0.01, and (C) 0.02. All *F*(*t*) curves are well described by a sum of two exponential decays with a long-time residual *F_R_*: *F*(*t*) = *FR* + *C*_1_ exp(−*t*/*τ*_1_) + *C*_2_ exp (−*t*/*τ*_2_) as indicated by the representative fit (solid black line) shown in (A). (D-H) The residual force *F_R_* (D), decay times *τ*_1_ (E) and *τ*_2_ (F), and corresponding fractional coefficients *c*_1_ = *C*_1_/(*C*_1_ + *C*_2_) (G) and *c*_2_ = *C*_2_/(*C*_1_ + *C*_2_) (H), are determined from the fits and plotted as functions of *R_α_* for *R_p_* = 0 (open squares, dotted connecting lines), *R_p_* = 0.01 (half-filled circles, dashed connecting lines), and *R_p_* = 0.02 (solid diamonds, solid connecting lines).

To understand the mechanisms underlying each exponential term, we turn to predicted relaxation timescales for entangled semiflexible polymers [55]. The fastest predicted relaxation mechanism is the mesh time *τ_mesh_*, i.e., the time for entangled polymers to feel the surrounding mesh [18,55,59], which depends on the network mesh size *ζ* and filament persistence length *l_p_* as *t_mesh_* ≃ 4*ζ*^4^/*l_p_* [18,55,63,64]. Assuming *ζ* = 0.3/*c*^1/2^ = 0.6 *μ*m for systems with and without phalloidin and considering the increased persistence length of phalloidin-stabilized filaments, the predicted *t_mesh_* values are 0.05 s and 0.03 s in the absence and presence of phalloidin, respectively. Our measured *τ*_1_ values for *R_p_* = 0, are comparable to the predicted mesh time, with an average value of *τ*_1_ (0.07 0.03) s and a modest increase with increasing *R_α_* (Fig 5E). This increase may be due to increasing mesh size as filaments begin to form bundles such that the effective concentration of fibers comprising the network is lower. Surprisingly, in contrast to predictions, the addition of phalloidin substantially increases *τ*_1_, with *R_α_*-averaged values of ⟨*τ*_1_⟩ ≃ (0.27 ± 0.14) s and (0.24 ± 0.04) s for *R_p_* = 0.01 and 0.02, respectively. Given the strong dependence of *t_mesh_* on the mesh size, these longer timescales may arise from the increasing mesh size of phalloidin-stabilized networks due to bundling, which reduces the effective concentration of distinct fibers comprising the network (as we see in Fig 2). Specifically, a <2-fold increase in *ζ* with *R_α_* would result in the ~8-fold longer timescales we measure compared to theoretical predictions (i.e. ~0.24 s versus ~0.03 s).

Unlike our measured *τ*_1_ values, *τ*_2_ appears largely independent of both *R_p_* and *R_α_* suggesting that the underlying relaxation mechanism is independent of filament stiffness, connectivity, or other properties of the network (Fig 5F. We also recall that *α*-actinin is a transient crosslinker, and its dissociation rate from actin has been shown to control the low-frequency relaxation of *α*-actinin-crosslinked actin networks in the linear regime [21]. These previous studies have reported dissociation-mediated relaxation timescales of ~1–2.5 s, with minimal dependence on *R_α_*. Our measured *τ*_2_ values, similarly insensitive to *R_α_*, are in close agreement with these previously reported values [21] with 〈*τ*_2_〉 = (1.94 ± 1.11) s, (2.19 ± 0.63) s, and (2.19 ± 1.20) s, for *R_p_* = 0, 0.01 and 0.02, respectively. We thus attribute our slow relaxation timescale to transient unbinding and rebinding of *α*-actinin crosslinks that allow for dissipative network rearrangement.

To shed further light on the impact of crosslinking and stabilization on the relaxation dynamics, we also evaluate the relative contributions of the fast and slow relaxation mechanisms (associated with *τ*_1_ and *τ*_2_, respectively) to the overall stress relaxation by evaluating the fractional coefficient of the corresponding exponential decaying term, *c*_1_ = *C*_1_/(*C*_1_ + *C*_2_) and *c*_2_ = *C*_2_/(*C*_1_ + *C*_2_). As shown in Fig 5(G, H), similar to the relaxation timescales, the coefficients are largely insensitive to the crosslinker density *R_α_*. We also observe that the fast relaxation contributes more to phalloidin-stabilized networks compared to *R_p_* = 0 networks, seen by larger *c*_1_ values in Fig 5(G), with crosslinker unbinding playing a lesser role (Fig 5(H)). This effect likely arises from the fact that the values of *τ*_1_ are larger for phalloidin-stabilized networks compared to *R_p_* = 0 networks, such that fast relaxation dynamics span more of the measured relaxation phase so they can contribute more to the relaxation. Moreover, as crosslinking is the predominant mechanism driving the viscoelastic response of non-stabilized networks, we expect the crosslinker dynamics to play an important role in the relaxation. Conversely, for phalloidin-stabilized networks, the increased filament stiffness and bundling also contribute substantially to the viscoelasticity, such that dissipative fluctuations on the scale of the mesh size (occurring over *τ_mesh_* ≃ *τ*_1_) contribute to the dynamics more strongly.

The complex effects of phalloidin stabilization on the mechanics of crosslinked actin networks suggest that similar stabilization may play an important role in the nonlinear response of entangled actin networks without crosslinkers. Namely, the impacts of filament stiffening and bundling, which rigidify the network while also increasing the mesh size and altering fiber connectivity, suggest that phalloidin-stabilization may depend strongly on the actin concentration, which dictates the network mesh size and degree of connectivity. To this end, we perform the same experimental protocols and analyses as described in the preceding sections for entangled actin networks (*R_α_* = 0) at five different actin concentrations ranging from 2x lower to 2x higher than *c_a_* = 5.8 *μ*M used for the crosslinked networks.

We find that the dependence of the force response on stabilization is surprisingly weak, in contrast to the crosslinked network results. This general finding suggests that it is the synergistic coupling of filament crosslinking and stiffening that leads to the pronounced effects of phalloidin stabilization seen in Figs 2–5. This insensitivity to stabilization may also indicate that crosslinking is necessary to facilitate substantial bundling of phalloidin-stabilized filaments that would otherwise remain as an entangled network of individual filaments. We explore these results and hypotheses below.

### Dependence of Nonlinear Force Response on Actin Concentration

We first evaluate the force response during strain for varying actin concentrations *c_a_* with *R_p_* = 0, 0.1 and 0.2 (Fig 6). As shown in Fig 6A,B and SI Fig S3, we find that the terminal force *F_t_* monotonically increases with *c_a_* for all networks, independent of *R_p_*. However, phalloidin-stabilization does modestly increase *F*(*x*) and its dependence on strain *x* compared to the *R_p_* = 0 case. Conversely, Fig 6C,D shows that the initial force *F*_0_ is higher for phalloidin-stabilized networks at low actin concentrations (*c_a_* ≤ 5.8 *μ*m) but transitions to being lower than *R_p_* = 0 networks at higher *c_a_*. This result suggests that at low *c_a_*, when the entanglement density is lower, and network connectivity is weaker, the stiffness of the filaments plays a principal role in the microscale force response. However, as the concentration, and thus the entanglement density and connectivity, increases, *F*_0_ for the *R_p_* = 0 networks becomes larger, indicating that it is the network connectivity that dominates over filament stiffness. Similar to our results for crosslinked networks (Fig 3), the lower initial force for the *R_p_* > 0 networks suggests the presence of weak bundling at high actin concentrations that decreases connectivity compared to the *R_p_* = 0 case.

**Figure 6.**
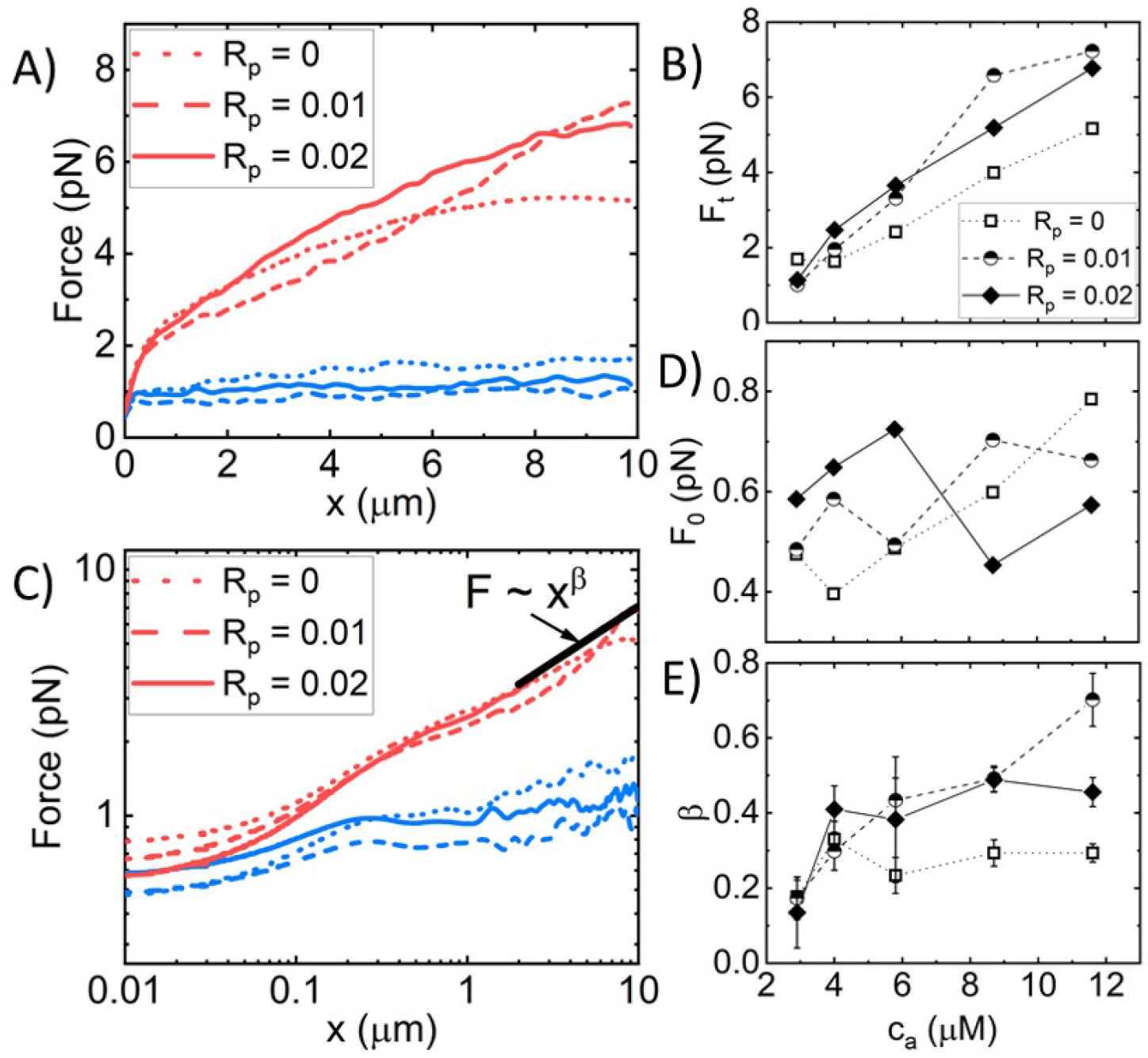
Effect of phalloidin stabilization and actin concentration on the nonlinear force response of entangled actin networks. (A) Force *F*(*x*) versus stage position *x* for actin networks subject to nonlinear straining (see Fig 1), for actin concentrations of *c_a_* = 2.9 *μ*m (blue) and *c_a_* = 11.6 *μ*m (red) and phalloidin:actin ratios of *R_p_* = 0 (dotted lines), *R_p_* = 0.01 (dashed lines), and *R_p_* = 0.02 (solid lines). (B) Terminal force reached at the end of the strain *F_t_* as a function of *c_a_* for *R_p_* = 0 (open squares, dotted connecting lines), *R_p_* = 0.01 (half-filled circles, dashed connecting lines), and *R_p_* = 0.02 (solid diamonds, solid connecting lines). (C-E) Data shown in (A) plotted on a log-log scale (C) to highlight the trends seen for the initial force *F*(*x* = 0) = *F*_0_ (plotted in D) and power-law scaling of *F*(*x*) near the end of the strain (plotted in E).

Also similar to crosslinked networks (Fig 3), the power-law scaling of *F*(*x*) at large lengthscales, i.e., *F*(*x*) ~ *x^β^*, is higher for phalloidin-stabilized networks compared to *R_p_* = 0 networks, indicating that filament stiffness plays the principal role in maintaining elastic storage at large lengthscales while crosslinks and entanglements can rearrange and dissipate induced stress over these spatiotemporal scales. However, while phalloidin-stabilization substantially increased the heterogeneity in the force response of crosslinked networks (Fig 3, SI Fig S1), suggestive of increased bundling, we observe no similar increase for entangled networks (SI Figs S2, S3). The phalloidin-mediated increase in *F_t_* and *β* is also weaker for entangled networks compared to crosslinked networks for all *c_a_* and *R_α_* values (Fig 6B,E), in line with our understanding that bundling of phalloidin-stabilized networks is facilitated by crosslinkers that bridge bundles.

The effect of phalloidin-stabilization on the stress stiffening and softening during strain, as well as the force relaxation following strain, is markedly weaker for entangled networks compared to crosslinked networks (Fig 3). While the initial and terminal stiffness *K*_0_ and *K_t_* are modestly larger for phalloidin stabilized networks, the degree of stiffening *K_max_*/*K*_0_ and the timescale over which the network yields to the terminal regime *t_y_* are largely insensitive to stabilization. Likewise, the relaxation timescales, *τ*_1_ and *τ*_2_, their relative contribution to the relaxation *c*_1_ and *c*_2_ and the residual force *F_R_* are all statistically indistinguishable across varying *R_p_* values (SI Fig S5(D–H)).

## 4. Conclusions

Networks of semiflexible actin filaments play critical roles in various cellular processes ranging from cell motility to stiffening to mechanosensation. Central to these mechanical processes are actin-binding proteins that crosslink and stabilize actin filaments. The structural changes that these ABPs mediate, in turn, tune the mechanical response of the networks to strain. Here, we used optical tweezers microrheology to elucidate the coupled effects of crosslinking and stabilization on the nonlinear force response and relaxation dynamics of entangled actin networks. Specifically, we study actin networks crosslinked by the transient crosslinker *α*-actinin at ABP:actin molar ratios of *R_α_* = 0 – 0.03, and stabilized by phalloidin at ABP:actin molar ratios of *R_p_* = 0, 0.01 and 0.02. Our results reveal complex relationships between crosslinking and stabilization that lead to emergent mechanical properties such as suppressed stress-stiffening and enhanced sustained elasticity. Notably, the effect of stabilization on the force response features of entangled actin networks is substantially weaker than for crosslinked networks, independent of the actin concentration. Taken together, our results demonstrate that crosslinking and phalloidin-mediated stiffening of actin filaments act synergistically to promote bundling while maintaining connectivity - both essential for soliciting a strong elastic response to nonlinear straining. However, at sufficiently high crosslinking, bundling comes at the cost of connectivity, leading to a nonmonotonic dependence on key mechanical properties such as softening time, terminal response elasticity, and microscale heterogeneity in force response. Beyond the relevance to cytoskeletal mechanics, our results may generally provide insight into the coupled roles of filament rigidity and connectivity on the nonlinear response of polymer networks and hydrogels.

## Supporting information

https://drive.google.com/drive/folders/18lMXQrECRzhRRNoL0nVif3UdAImtu0SV

## Data Availability Statement

The data presented in this study are available on request from the corresponding author.

## Conflicts of Interest

“The authors declare no conflict of interest.”

## Funding

This research was funded by a WM Keck Research Grant and NSF DMREF-2119663 awarded to RMRA, and startup funding from Bucknell University awarded to BJG.

## Notes

### Competing Interest Statement

The authors have declared no competing interest.

