## Supplementary material for "Nonlinear microscale mechanics of actin networks governed by coupling of filament crosslinking and stabilization": https://drive.google.com/drive/folders/18lMXQrECRzhRRNoL0nVif3UdAImtu0SV

*Article*

### 1. Force response of the individual trials during the strain phase

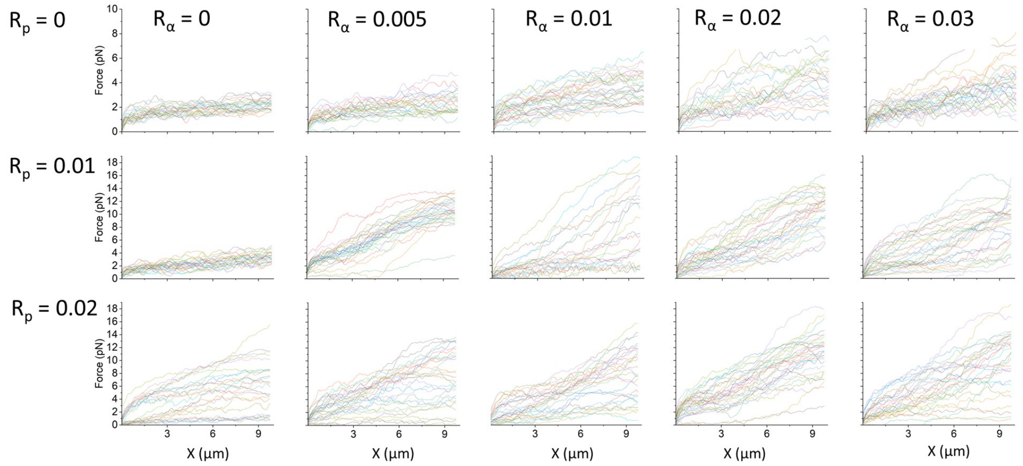


**Figure S1.** $R_\alpha$-actinin dependent elastic response of actin networks to nonlinear straining for the 30 different trials measured with 30 different probe particles by varying the molar ratio of $R_\alpha - actinin$:actin and $R_p$ as indicated on the top and left hand side.

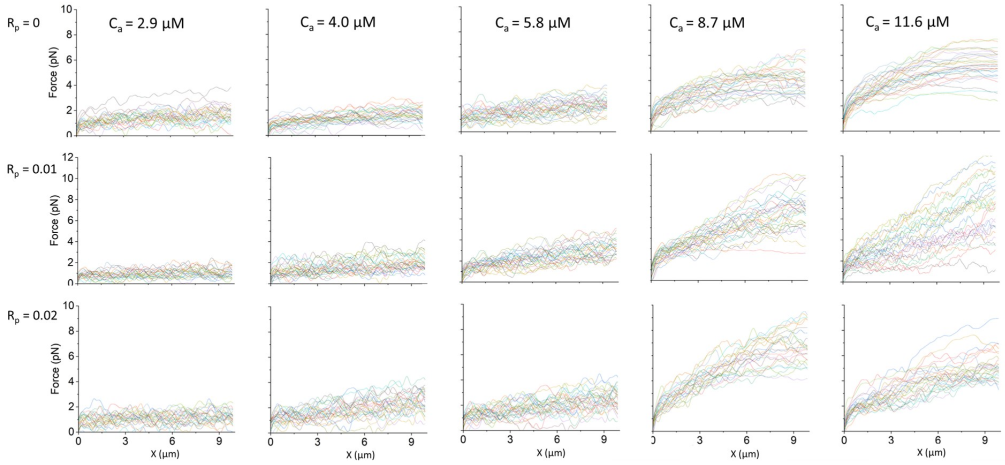


**Figure S2.** Concentration dependent elastic response of actin networks to nonlinear straining for the 30 different trials measured with 30 different probe particles by varying the actin concentration $c_a$ and the molar ratio of phalloidin:actin $R_p$ on the top and left hand side.

check for updates

**Citation:** Dwyer, M.; Robertson-Anderson, R.; Gurmessa, B. . *Preprints* **2022**, *1*, 0. https://doi.org/

**Publisher's Note:** MDPI stays neutral with regard to jurisdictional claims in published maps and institutional affiliations.

**Copyright:** © 2022 by the authors. Licensee MDPI, Basel, Switzerland. This article is an open access article distributed under the terms and conditions of the Creative Commons Attribution (CC BY) license (https://creativecommons.org/licenses/by/4.0/).

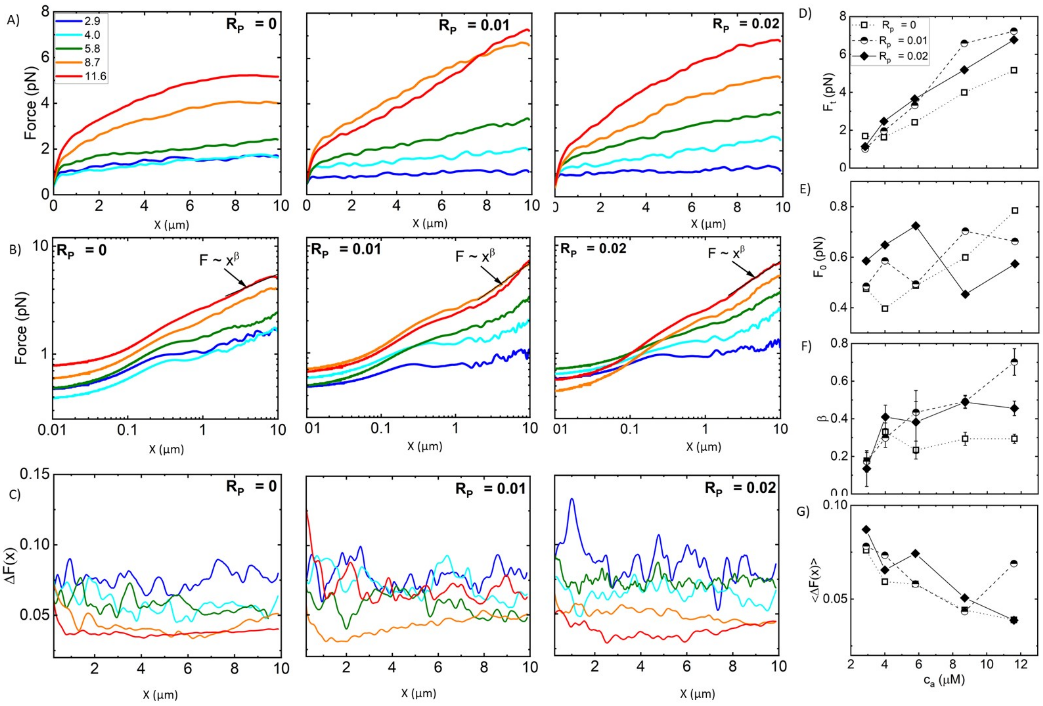


**Figure S3.** Concentration dependent elastic response of actin networks to nonlinear straining. (A) Average force $F(x)$ (shown in SI Fig. S2) versus stage position $x$ measured for actin networks subject to nonlinear straining. Different curves in each panel correspond to actin concentration $c_a$ = 0 - 0.03, color-coded according to the legend in A. Different panels display data for phalloidin:actin molar ratios of $R_p$ = 0 (left), $R_p$ = 0.01 (middle), and $R_p$ = 0.02 (right). (B) Data shown in (A) plotted on a log-log scale to highlight the trends seen for the initial force $F(x = 0) = F_0$ and power-law scaling of $F(x)$ near the end of the strain ($x \approx 1$). Fitting the large strain data to a power-law F(x)$\sim x^\beta$ yields the scaling exponent $\beta$ that quantifies the degree of elastic storage maintained at the end of the strain. (C) Fractional spread in force $\Delta F(x)$ for each position $x$ and each condition, determined by computing the standard error across 30 individual trials and normalizing by the average value plotted in A: $\Delta F(x) = SEF(x)/\langle F(x)\rangle$. (D-G) Metrics computed from the data shown in A-C as a function of $c_a$ for $R_p$=0 (open squares, dotted connecting lines), $R_p$=0.01 (half-filled circles, dashed connecting lines), and $R_p$=0.02 (solid diamonds, solid connecting lines): (D) Terminal force reached at the end of the strain $F_t$; (E) Initial force measured at the beginning of the strain $F_0$; (F) Power-law scaling exponent $\beta$ determined from fits to F(x)$\sim x^\beta$ depicted in B; (G) $\Delta F(x)$ averaged over the strain position $x$, resulting in a single value for each curve shown in (C) with error bars denoting the standard error across $x$ values.

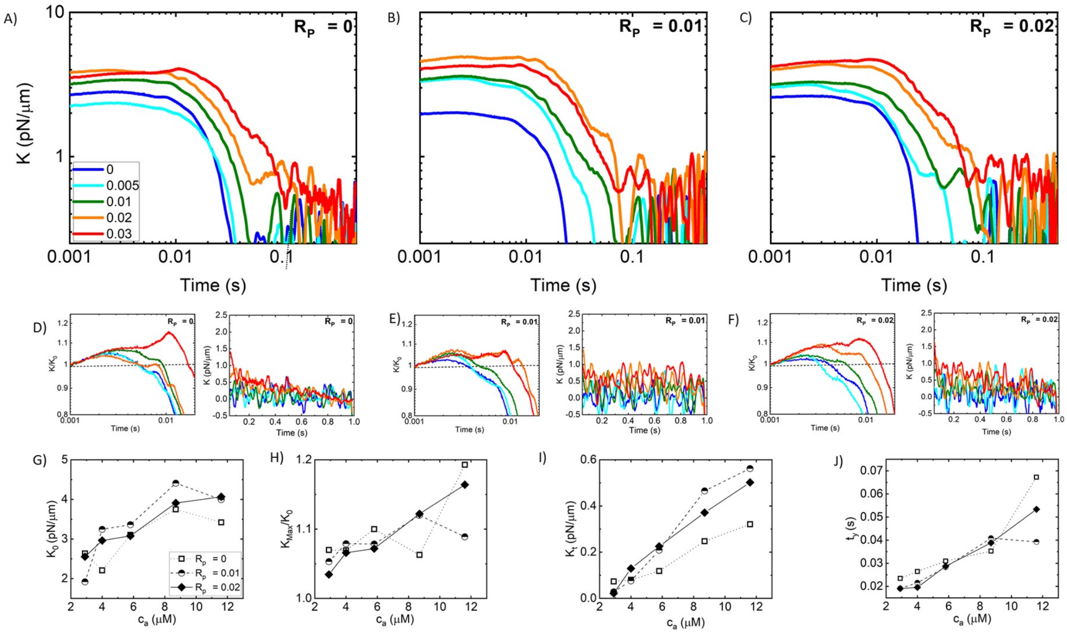


**Figure S4.** Effective differential modulus $K(t) = dK(x,t)/dx$ as a function of time during strain $t = v/x$, computed from the data shown in SI Fig. S3A. The color-coded curves in each panel are for $R_\alpha$ values indicated in the legend and the different panels show data for (A) $R_p$ = 0, (B) $R_p$ = 0.01, and (C) $R_p$ = 0.02. (D-F) $K(t)$ data shown in (A-C) normalized by the corresponding initial value $K_0$ are shown in the left panels. The horizontal dashed line at $K(t)/K_0$=1 guides the eye to show stress stiffening ($K(t)/K_0 > 1$) or softening ($K(t)/K_0 < 1$). Right panels show zoom-ins of $K(t)$ near the end of the strain where $K(t)$ is approximately constant. (G-J) Metrics computed from the data shown in A-C as a function of $c_a$ for $R_p$=0 (open squares, dotted connecting lines), $R_p$=0.01 (half-filled circles, dashed connecting lines), and $R_p$=0.02 (solid diamonds, solid connecting lines): (G) Initial differential modulus $K_0$, (H) Degree of stress-stiffening, quantified as $K_{max}/K_0$ and equal to 1 for networks which only display softening (i.e.$K_{max} = K_0$), (I) Terminal stiffness $K_t$ computed by averaging over the $K(t)$ data shown in the right-hand panels of D-F, (J) Softening time $t_y$, defined as the time at which $K(t) = K_0/e$

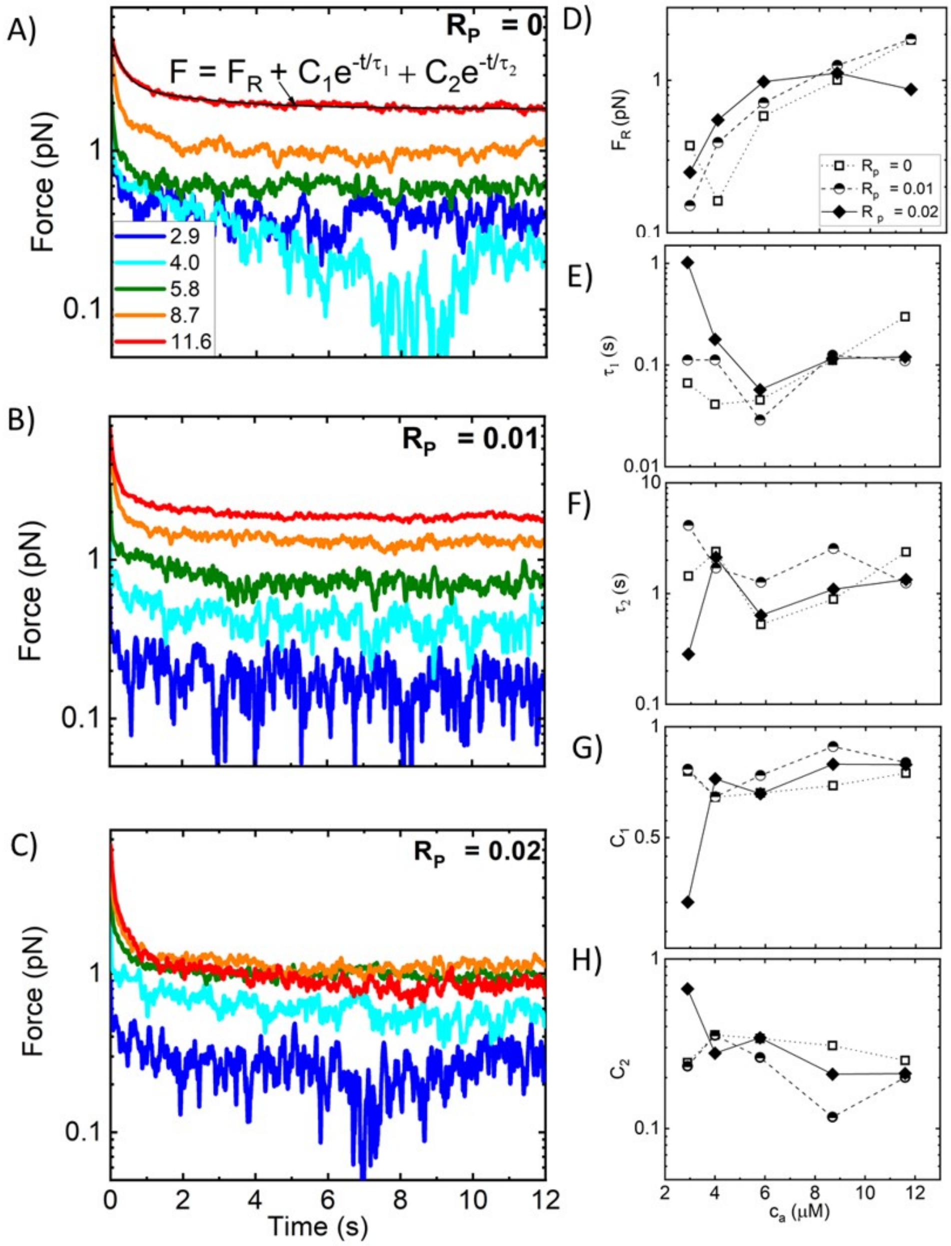


**Figure S5. Strain-induced force exponentially relaxes over time to a residual value $F_R$ dependent on $R_p$ and $c_a$.** (A-C) Time-dependent relaxation of force $F(t)$ following strain for actin networks with varying $c_a$ values, color-coded according to the legend, and with $R_p$ values of (A) 0, (B) 0.01, and (C) 0.02. , $R_p$ = 0.02. All $F(t)$ curves are well described by a sum of two exponential decays with a long-time residual $F_R$: $F(t) = F_R + C_1 \exp(-t/\tau_1) + C_2 \exp(-t/\tau_2)$ as indicated by the representative fit (solid black line) shown in (A). (D-H) The residual force $F_R$ (D), decay times $\tau_1$ (E) and $\tau_2$ (F), and corresponding fractional coefficients $c_1 = C_1/(C_1 + C_2)$ (G) and $c_2 = C_2/(C_1 + C_2)$ (H), are determined from the fits and plotted as functions of $c_a$ for $R_p$=0 (open squares, dotted connecting lines), $R_p$=0.01 (half-filled circles, dashed connecting lines), and $R_p$=0.02 (solid diamonds, solid connecting lines).
